# Single cell RNA sequencing reveals a novel, microglia-like cell type in cerebrospinal fluid during virologically suppressed HIV

**DOI:** 10.1101/310128

**Authors:** Shelli F. Farhadian, Sameet S. Mehta, Chrysoula Zografou, Jenna Pappalardo, Jennifer Chiarella, David A. Hafler, Serena S. Spudich

**Affiliations:** Department of Medicine, Section of Infectious Diseases, Yale School of Medicine, New Haven, CT, USA.; Yale Center for Genome Analysis, Yale School of Medicine, New Haven, CT, USA.; Department of Neurology, Yale School of Medicine, New Haven, CT, USA.; Department of Immunobiology, Yale School of Medicine, New Haven, CT, USA.

## Abstract

Central nervous system (CNS) immune activation in an important driver of neuronal injury during several neurodegenerative and neuroinflammatory diseases. During HIV infection, CNS immune activation is associated with high rates of neurocognitive impairment, even with sustained long-term suppressive antiretroviral therapy (ART). However, the cellular subsets that drive immune activation and neuronal damage in the CNS during HIV infection and neurological conditions remain unknown, in part because CNS cells are difficult to access in living humans. Using single cell RNA sequencing (scRNA-seq) on cerebrospinal fluid (CSF) and blood from adults with HIV, we identified a rare (<5% of cells) subset of myeloid cells in CSF presenting a gene expression signature consistent with neurodegenerative disease associated microglia. This highlights the power of scRNA-seq of CSF to identify rare CNS immune cell subsets that may perpetuate neuronal injury during HIV infection and other conditions.

## Introduction

Neuronal injury during infections and other inflammatory processes may be due to both the direct effect of a toxic pathogen as well as bystander effect of nearby, activated immune cells. During HIV infection, CNS immune activation has been implicated in the persistently high rates of neurological impairment seen in adults with HIV, a phenomenon that persists even during virological suppression with ART[1–3]. However, the specific cellular subsets and genes that drive CNS immune activation during HIV and other neuroinflammatory conditions remain incompletely understood. This is in part because brain tissue is largely inaccessible for routine studies of neurological disease. New tools are thus needed to interrogate CNS cells in living humans.

Cerebrospinal fluid is produced within the brain in the choroid plexus and we have previously shown in other neuroinflammatory disease that CSF immune cells reflect infiltrating brain parenchymal immune cells[4,5]. Thus, the CSF can provide unique diagnostic information in infectious, inflammatory and neurodegenerative CNS disorders[6–8]. In adults with HIV infection, while rare studies have analyzed brain tissue obtained at autopsy of affected patients to understand the cellular basis for persistent CNS immune activation during HIV infection, a majority of studies have used CSF analysis, though primarily focusing on soluble biomarkers that reflect non-specific immune activation, or on flow cytometry analysis of CSF immune cells[9–12]. However, a major limitation of flow cytometry is that cell populations are identified based only on a small set of surface markers, thus potentially missing rare and important cells that are unique to the CNS and which may be transcriptionally distinct but may not display different surface markers.

Myeloid-lineage cells in the CNS, including microglia and circulating monocytes, are particularly challenging to study using conventional approaches. These cells have been proposed to play a role in causing or exacerbating neuronal injury during HIV infection. During acute infection, monocytes traffic to the CNS where they have been proposed to contribute to neuroinflammation and may contribute to establishment of latent CNS infection[13]. Microglia, the resident tissue macrophages of the CNS, are a potential site for persistent latent infection or low-level viral replication in the CNS, as well as a potential driver of neuronal injury, even during ART[14,15]. CSF studies in adults with virologically suppressed HIV have revealed elevated levels of neopterin and soluble CD163, biomarkers that are associated with monocyte and microglial activation, further supporting a role for monocytes and microglia in perpetuating neuronal injury during HIV infection[9,16]. However, since myeloid cells comprise a minor population of CSF cells (<10%), they are challenging to detect and fully characterize through cytometry based studies[9]. Moreover, microglia have thusfar only been characterized in humans through autopsy brain tissue or through *in vitro* cell-culture based studies.

Here we use massively parallel single-cell RNA-seq to characterize the immune cell landscape of CSF in HIV-infected individuals with virologic suppression. Recent advances in scRNA-seq now allow for simultaneous examination of >10,000 single cell transcriptomes and have led to the characterization of new immune cell subsets when applied to whole blood as well as various human tissues in both healthy and diseased states[17–19]. We performed scRNA-seq on large volume CSF and paired blood samples from individuals with virologically suppressed HIV to discover cell populations that are enriched in the CSF of individuals with HIV on ART, including myeloid cell subsets. By studying CSF immune cells in an unbiased, surface-marker-free approach, we sought to identify *de novo* cell populations that are associated with CNS immune activation.

## Results and Discussion

High-quality cDNA was produced from single-cell whole transcriptomes derived from CSF and blood that were collected in parallel **(Figure 1A)**.

**Figure 1.**
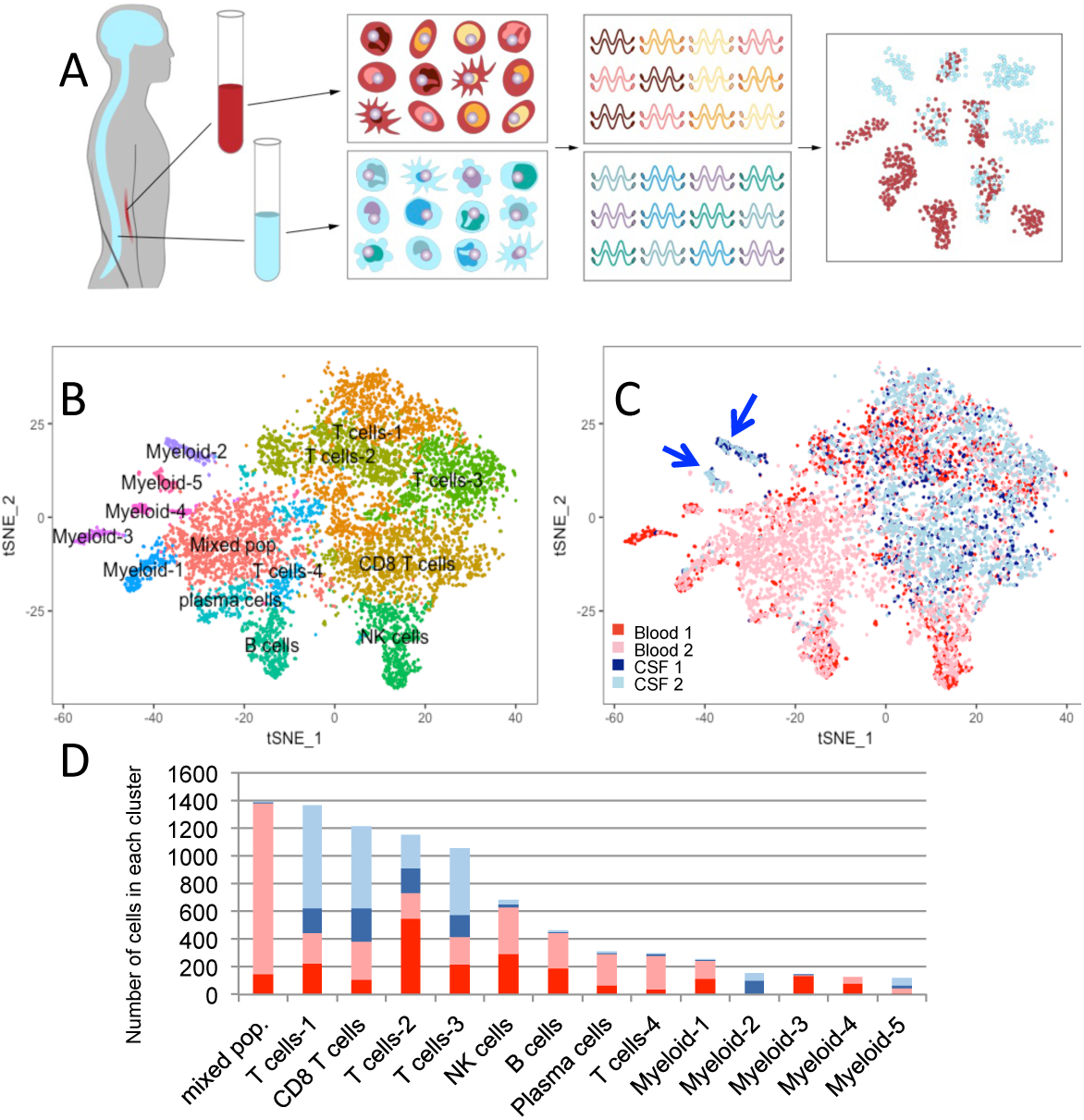
Single cell RNA sequencing of paired blood and CSF samples. **1A:** CSF and blood were collected and processed in parallel. Single PBMCs and CSF cells were applied to separate arrays, with 10,000 cells applied to each SeqWell array. Single cell whole transcriptomes were tagged to retain information about the cell, and then sequenced. Sequencing data from single cells was pooled for cluster analysis based on differential gene expression, revealing clusters that are comprised predominantly of CSF cells. **1B:** tSNE visualization of cell populations identified through unsupervised cluster analysis of blood and CSF derived from HIV+ individuals. Clusters are labeled based on expression of canonical marker genes (see also Supplemental Figure 1). **1C:** tSNE visualization of cells, labeled by sample and tissue of origin, revealing two myeloid clusters that are predominantly comprised of CSF cells (blue arrows). **1D** Bar plots quantifying the number of cells from blood and CSF from each sample in each cluster.

We detected a mean of 775 genes per single cell in CSF and 758 genes per cell in blood. CSF and peripheral blood mononuclear cell (PBMC) single-cell transcriptomes contained similar proportions of mitochondrial genes (mean 10%). A total of 8,774 single cell transcriptomes derived from CSF (3,160 cells) and PBMCs (5,580 cells) of two HIV+ participants were pooled for initial unsupervised cluster analysis that did not rely on known markers of cell types, thus allowing for the identification of novel cell populations. Fourteen clusters representing distinct immune cell subsets were identified **(Figure 1B**). Identities were assigned to cell clusters based on expression levels of canonical marker genes and of differentially expressed genes (Supplementary Figure 1). We then overlayed the tissue of origin of original of each cell (i.e., blood or CSF), and in this way, identified differences in cellular distributions between blood and CSF **(Figure 1C)**. We found a majority of cells in CSF were T lymphocytes (89%), with smaller populations of myeloid cells (7%), NK cells (<2%), and B cells and plasma cells (<1%). When compared to blood, CSF contained higher proportions of CD8 T cells, and two myeloid cellular subsets **(Figure 1D)**.

We next performed sub-analysis of the myeloid cells identified from the larger, combined sample of blood and CSF cells **(Figure 2A)**. We found five distinct myeloid cellular subsets, two of which consisted of cells found predominantly (>50% of cells) in CSF (Myeloid-2 and Myeloid-5). Differential gene expression analysis revealed that Myeloid-5 consists of cells with high levels of expression of genes found in myeloid-derived dendritic cells, including *CD1C* and *FCER1A*.

**Figure 2.**
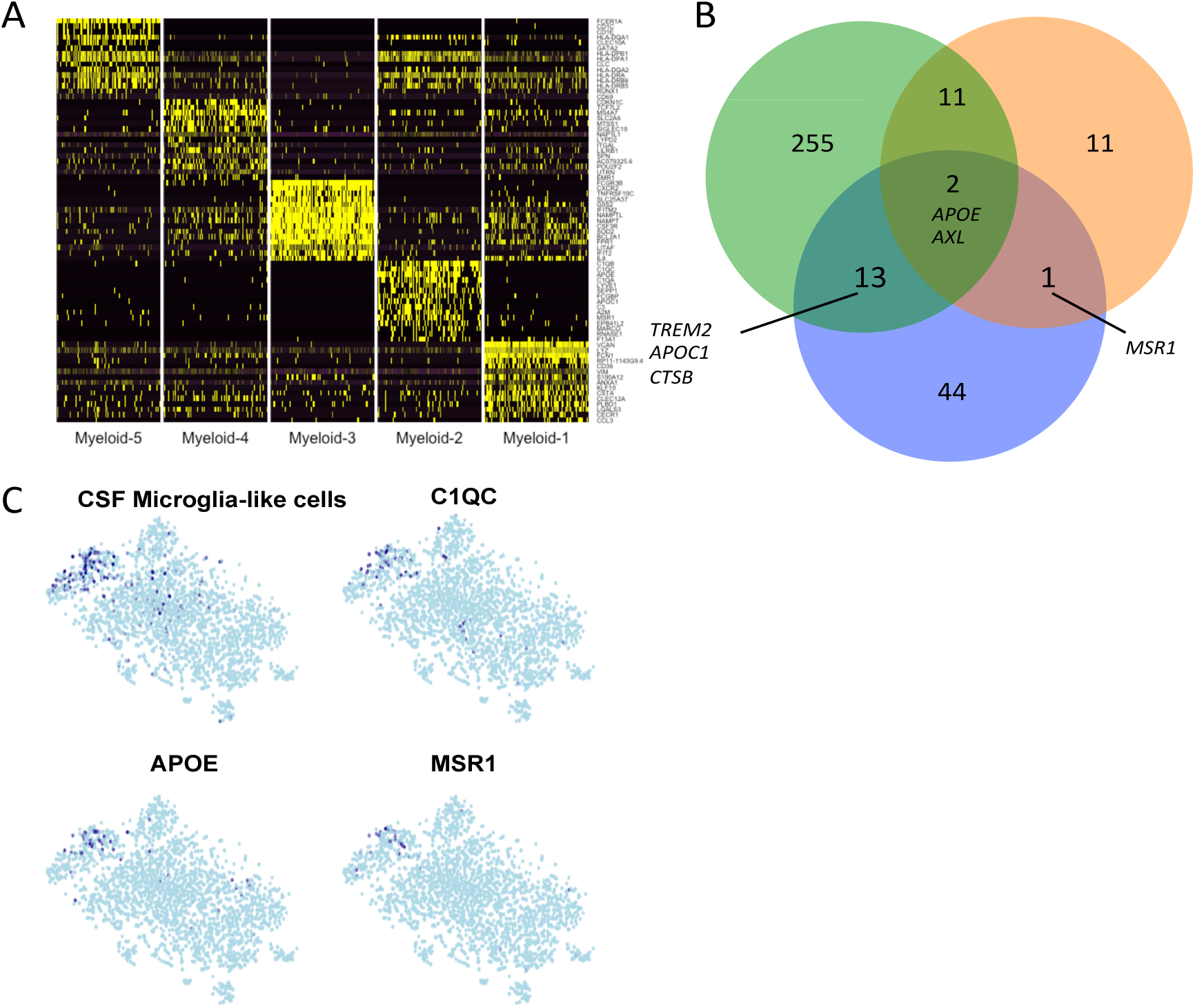
Single cell RNA sequencing reveals a disease associated microglia-like cellular subset in cerebrospinal fluid. **2A:** Heatmap showing the top fifteen genes that most differentiate each of the five myeloid subsets. **2B:** Venn diagram shows genes that were found in Myeloid-2 (blue) and across two independent transcriptomic studies of neurodegenerative disease-associated microglial cells derived from mouse brain (green[17] and orange[22]). **2C:** Examination of an independent CSF sample from an HIV+ participant reveals a group of cells with co-expression of the genes that characterize the CSF microglia-like myeloid subset.

Myeloid-2 is a myeloid subset comprised almost exclusively of CSF cells and is characterized by high expression of 60 distinct, non-ribosomal genes when compared to all other myeloid subsets (FDR < 0.1) (Supplementary Table 1). Several of these genes are produced in CNS almost exclusively by microglia, including *C1QA-C* and *TREM2* [20,21].

We therefore compared the genes that characterize Myeloid-2 to recently published transcriptomic studies that examined microglia derived from mouse models of neurodegenerative diseases[17,22] **(Figure 2B)**. We found significant overlap between genes that are overexpressed in Myeloid-2 and genes that are enriched in neurodegenerative-disease associated microglia, including *APOE, AXL*, and *TREM2*. Furthermore, we found that, compared to the four other myeloid subsets we identified, Myeloid-2 expressed higher levels of *CTSB, APOC1*, and *MSR1(*CD204), molecules that are also associated with the microglial neurodegenerative phenotype[22]. We therefore reason that Myeloid-2 consists of CSF neurodegenerative-disease associated microglia-like cells.

We asked whether this subset of CSF microglia-like cells could be detected in CSF in an independent sample derived from an HIV+ participant. We found a group of 93 cells with high aggregate expression of the set of genes that defined the “Myeloid-2” population of CSF-associated microglia-like cells **(Figure 2C)**. This represents 3.3% of all cells in the independent CSF sample. This indicates that the rare, CSF-associated microglia-like cells we identified earlier are also present in CSF from an independent HIV+ participant.

Thus, our study used an unbiased, surface marker-free approach to characterize CNS immune cell populations during virologically suppressed HIV infection and identified microglia-like cells in CSF. To our knowledge, this is the first study to characterize the full immune cell landscape in CSF at the transcriptional level, and the first to identify circulating microglia-like cells in CSF. These CSF-associated microglia-like cells are rare, representing <5% of all cells we analyzed in CSF, and thus would likely not be reliably detected using a traditional flow-cytometry based studies of CSF.

CNS myeloid cells, and microglia in particular, have been proposed to play several important roles in HIV infection, including acting as a potential site of CNS viral replication. In their normal, homeostatic state, microglia play an important role in maintaining neuronal integrity, by promoting immune responses to infection and through clearance of debris and plaque. During chronic neurological diseases, including Alzheimer’s disease and ALS, microglia may become dysfunctional and promote neuronal injury. This switch, from homeostatic to neurodegenerative microglia, has been linked to an APOE-TREM2 signaling pathway that disrupts the normal microglial response to damaged neurons and other debris. Likewise, microglial dysregulation has been proposed as a possible mechanism for neuronal toxicity during HIV infection, and our results suggest a potential role for the APOE-TREM2 pathway in mediating neurodegeneration during HIV.

In conclusion, our study identifies microglia-like cells in human CSF with a transcriptional profile that is similar to disease-associated microglia in animal models of neurodegenerative disease. This demonstrates a potential mechanistic link between pathways of neuronal injury in HIV and other neurodegenerative conditions. Furthermore, our study illustrates the potential for genomic-based studies of CSF to uncover rare immune cell populations that may drive a wide-range of CNS disease.

## Methods

### Study participants and procedures

Research participants were seen for study visits at the Yale School of Medicine. Participants were HIV+ and on stable ART with plasma HIV RNA levels <20 copies/mL for >1 year. None had active neurological disease or other infection. Participants underwent large volume lumbar puncture (25cc CSF removed) and blood draw. Blood and CSF were processed separately for single cell RNA sequencing using SeqWell[19].

### CSF and blood processing

Fresh CSF and blood were processed within one hour of collection. CSF was centrifuged at 1500 rpm for 10 minutes, and the isolated cells were immediately applied to SeqWell arrays. Peripheral blood mononuclear cells (PBMCs) were isolated via Ficoll gradient. Approximately 10,000 cells were loaded onto each SeqWell array, with one array per CSF or PBMC sample. Cells were processed as in *Geirahn et al*[19]. Briefly, cells were added to custom microwell arrays that were pre-loaded with “Drop-Seq beads” (Chemgenes CSO-2011). These beads contain oligonucleotides comprised of a unique cellular barcode, poly-T for mRNA capture, a unique molecular identifier, and a PCR handle[18]. Each microwell thus ideally contained one barcoded bead and one cell. A semipermeable membrane was applied to each array, cells were lysed, and mRNA was hybridized to the barcoded beads. Membranes were then removed; microwell contents were pooled; cDNA was generated; and Nextera sequencing libraries were prepared. Sequencing was performed on the Illumina HiSeq 4000 platform at approximately 50,000 reads per cell.

### Single cell transcriptome analysis

Low quality cells containing <500 or >2500 genes detected were removed, as well as the 5% of cells with highest mitochondrial content. Genes that were present in less than 3 cells were excluded from analysis. Gene expression values were then normalized, scaled, and log-transformed.

The detection of highly variable genes and unbiased identification of cell clusters was performed with Seurat[23]. Single cell transcriptomes from CSF and blood from two HIV+ participants were pooled prior to unsupervised cluster analysis. Four clusters that were characterized primarily by mitochondrial and ribosomal gene transcripts (indicating low quality cells) were removed. Histograms were generated to determine the relative contributions of CSF and of blood cells to each cluster, thus permitting the identification of clusters that were comprised predominantly of CSF cells. For analysis of the independent HIV+ CSF sample (Figure 2C), we performed supervised analysis to look for cells that contained the transcripts that were previously identified as marker genes for “Myeloid-2,” the microglia-like subset identified during the unsupervised analysis.

### Study approval

This study was approved by the Yale University Human Investigations Committee. Informed consent was obtained using protocols approved by the Yale Human Research Protection Program.

## Author contributions

SFF and SS designed the study. SF and CZ conducted SeqWell experiments. SSM and SFF performed data analysis. JP contributed to data analysis. JC contributed to participant recruitment. DM provided reagents and contributed to study design. SFF and SS wrote the manuscript.

## Acknowledgements

We thank the participants who donated samples for this study; T. Kirchwey, M. Chintanaphol, and L. Le; and Todd Gierahn, Brittany Goods, and the Christopher Love laboratory/Koch Institute for Integrative Cancer Research at MIT for providing arrays and technical assistance. SFF was supported by NIH T32AG1934.

## References

1. Tozzi V, Balestra P, Bellagamba R, Corpolongo A, Salvatori MF, Visco-Comandini U, et al. Persistence of neuropsychologic deficits despite long-term highly active antiretroviral therapy in patients with HIV-related neurocognitive impairment: prevalence and risk factors. J. Acquir. Immune Defic. Syndr. 2007;45:174–82.

2. Heaton RK, Franklin DR, Deutsch R, Letendre S, Ellis RJ, Casaletto K, et al. Neurocognitive change in the era of HIV combination antiretroviral therapy: The longitudinal CHARTER study. Clin. Infect. Dis. 2015;60:473–80.

3. Grauer OM, Reichelt D, Grüneberg U, Lohmann H, Schneider-Hohendorf T, Schulte-Mecklenbeck A, et al. Neurocognitive decline in HIV patients is associated with ongoing T-cell activation in the cerebrospinal fluid. Ann. Clin. Transl. Neurol. 2015;2:906–19.

4. Lovato L, Willis SN, Rodig SJ, Caron T, Almendinger SE, Howell OW, et al. Related B cell clones populate the meninges and parenchyma of patients with multiple sclerosis. Brain. 2011;134:534–41.

5. Stern JNH, Yaari G, Vander Heiden JA, Church G, Donahue WF, Hintzen RQ, et al. B cells populating the multiple sclerosis brain mature in the draining cervical lymph nodes. Sci. Transl. Med. 2014;6.

6. Iorio R, Lennon VA. Neural antigen-specific autoimmune disorders. Immunol. Rev. 2012;248:104–21.

7. Damkier HH, Brown PD, Praetorius J. Cerebrospinal fluid secretion by the choroid plexus. Physiol. Rev. 2013;93:1847–92.

8. Tyler KL. Herpes simplex virus infections of the central nervous system: encephalitis and meningitis, including Mollaret’s. Clin. Infect. Dis. 2004;11 Suppl 2:57–64.

9. Ho EL, Ronquillo R, Altmeppen H, Spudich SS, Price RW, Sinclair E. Cellular Composition of Cerebrospinal Fluid in HIV-1 Infected and Uninfected Subjects. PLoS One. 2013;8.

10. Yilmaz A, Yiannoutsos CT, Fuchs D, Price RW, Crozier K, Hagberg L, et al. Cerebrospinal fluid neopterin decay characteristics after initiation of antiretroviral therapy. J. Neuroinflammation. 2013;10:62.

11. Edén A, Marcotte TD, Heaton RK, Nilsson S, Zetterberg H, Fuchs D, et al. Increased Intrathecal Immune Activation in Virally Suppressed HIV-1 Infected Patients with Neurocognitive Impairment. PLoS One. 2016;11.

12. Burdo TH, Weiffenbach A, Woods SP, Letendre S, Ellis RJ, Williams KC. Elevated sCD163 in plasma but not cerebrospinal fluid is a marker of neurocognitive impairment in HIV infection. AIDS. 2013;27:1387–95.

13. Williams D, Veenstra M, Gaskill P, Morgello S, Calderon T, Berman J. Monocytes Mediate HIV Neuropathogenesis: Mechanisms that Contribute to HIV Associated Neurocognitive Disorders. Curr. HIV Res. 2014;12:85–96.

14. Williams K, Burdo TH. Monocyte mobilization, activation markers, and unique macrophage populations in the brain: Observations from SIV Infected monkeys are informative with regard to pathogenic mechanisms of HIV infection in humans. J. Neuroimmune Pharmacol. 2012. p. 363–71.

15. Honeycutt JB, Wahl A, Baker C, Spagnuolo RA, Foster J, Zakharova O, et al. Macrophages sustain HIV replication in vivo independently of T cells. J. Clin. Invest. 2016;126:1353–66.

16. Neuenburg JK, Cho T a, Nilsson A, Bredt BM, Hebert SJ, Grant RM, et al. T-cell activation and memory phenotypes in cerebrospinal fluid during HIV infection. J. Acquir. Immune Defic. Syndr. 2005;39:16–22.

17. Keren-Shaul H, Spinrad A, Weiner A, Matcovitch-Natan O, Dvir-Szternfeld R, Ulland TK, et al. A Unique Microglia Type Associated with Restricting Development of Alzheimer’s Disease. Cell. 2017;169:1276–1290.e17.

18. Macosko EZ, Basu A, Satija R, Nemesh J, Shekhar K, Goldman M, et al. Highly parallel genome-wide expression profiling of individual cells using nanoliter droplets. Cell. 2015;161:1202–14.

19. Gierahn TM, Wadsworth MH, Hughes TK, Bryson BD, Butler A, Satija R, et al. Seq-Well: portable, low-cost RNA sequencing of single cells at high throughput. Nat. Methods. 2017;

20. Fonseca MI, Chu SH, Hernandez MX, Fang MJ, Modarresi L, Selvan P, et al. Cell-specific deletion of C1qa identifies microglia as the dominant source of C1q in mouse brain. J. Neuroinflammation. 2017;14.

21. Butovsky O, Jedrychowski MP, Moore CS, Cialic R, Lanser AJ, Gabriely G, et al. Identification of a unique TGF-β-dependent molecular and functional signature in microglia. Nat. Neurosci. 2014;17:131–43.

22. Krasemann S, Madore C, Cialic R, Baufeld C, Calcagno N, El Fatimy R, et al. The TREM2-APOE Pathway Drives the Transcriptional Phenotype of Dysfunctional Microglia in Neurodegenerative Diseases. Immunity. 2017;47:566–581.e9.

23. Satija R, Farrell JA, Gennert D, Schier AF, Regev A. Spatial reconstruction of single-cell gene expression data. Nat. Biotechnol. 2015;33:495–502.

